# Testing bottom-up cuing effects on target detection and discrimination in Bumblebees

**DOI:** 10.1101/2025.04.23.650250

**Authors:** Théo Robert, Marion Callendret, Chloe Sowels, Vivek Nityananda

## Abstract

Attention in vertebrates helps prioritise the processing of important sensory information and filter out irrelevant signals. The capture of attention by sudden or salient stimuli typically called bottom-up attention. Little is known about similar attentional process in insects, although they should be advantageous for insects as well. We therefore adapted two paradigms used to investigate bottom-up attention in primates to investigate it in bumblebees: a target detection task and a target discrimination task. For both tasks, we trained bees to choose between two locations on each side of a computer screen and collect a reward bellow a full contrast target displayed on the screen. During detection task tests, the contrast of the target was varied and it could be preceded by a cue flashed on the side of the target, the opposite side of the screen or not flashed at all. The discrimination task tests were similar but with a full contrast target on one side and a variable contrast distractor on the opposite side of the screen. We tested if the presence of the flash influenced the orientation and choices of the bees as well as their contrast sensitivity as has been seen in primates. Our results show no effect of the prior cue, suggesting that other paradigms might prove more useful to test these processes in insects.

## Introduction

Animals are constantly exposed to a multitude of sensory stimuli in their environment. The limited neural resources available to an animal means that these stimuli have to compete for processing and stimuli that are relevant to survival need to be prioritized. Vertebrates have therefore evolved attentional processes through which can prioritise the treatment of the relevant sensory stimuli and filter out noise (reviewed in Carrasco, 2011).

One form of attentional prioritization is based on spatial location. Humans are faster to detect a visual target at a particular location, if they are previously cued to that location (Posner, 1980) Conversely, the detection time increases if the subjects re cued towards a different location. Two distinct mechanisms underlie spatial attention. The first involves tuning sensory systems to increase the saliency of specific stimuli previously associated with a reward (Maunsell & Treue, 2006; Scolari et al., 2014). Primates can willingly increase their sensory sensitivity in specific spatial areas based on available information (Fernández et al., 2022). This active process, generally referred as top-down or endogenous attention, can help animals better detect relevant goals in the environment around them. In the second mechanism, subjects’ attention can be captured and directed towards a region in space by salient stimuli such a flash or a loud sound through an involuntary process called bottom-up or exogenous attention (Henderson & Macquistan, 1993).

Bottom-up attention has been demonstrated in many vertebrate taxa. In addition to humans, other - non-human - primates have also been shown to have such an attentional process (Bowman et al., 1993; Wang et al., 2015). Attentional processes in primates are thought to be supported by neural pathways in their neocortex (Bowling et al., 2020; Meyer et al., 2018; for a review see Behrmann et al., 2004). Similar attentional processes have, however, also been shown in birds (Quest et al., 2022; Shimp & Friedrich, 1993; Sridharan et al., 2014) and possibly in fish (Gabay et al., 2013), despite the lack of a neocortex. It therefore seems likely that this is an important cognitive feature that is evolutionarily selected for. We could therefore expect similar attentional processes to also have evolved in invertebrates.

One of the most noticeable effects of visual bottom-up attention in primates is a localised increase in contrast sensitivity. Cuing a subject’s attention towards a location allows them to perceive a subsequent target at that location at a lower contrast threshold, compared to when their attention is cued to another location (Barbot et al., 2011; Cameron et al., 2002; Fernández et al., 2019; Herrmann et al., 2010; Jigo & Carrasco, 2020). An increase in contrast sensitivity in response to a sudden localised change would be useful for an animal to better perceive and identify the cause of this change. It could, for example, allow the animal to quickly recognise a predator or rapidly identify potential prey.

Such adaptations would also be useful for insects. Yet very few studies have directly investigated attentional processes in these animals (reviewed in Nityananda, 2016). Weiderman and O’Carroll (2013) demonstrated that a visual neuron of a dragonfly (CSTMD1) responded selectively to one of two targets moving vertically at two different places in the visual field. More recent work showed that if one of the two locations was primed before the simultaneous presentation of the targets, the neuron was more likely to respond to the stimulus shown at the primed location (Lancer et al., 2019). Cuing effects on bottom-up attention in insects have also been shown behaviourally (Sareen et al 2011) in the fruit fly *Drosophila melanogaster*. In this study, the authors displayed two vertical bars on a circular screen to tethered flies. When the bars moved in opposite directions on the screen, the flies had an equal probability of turning in the direction of either bar. However, if one of the stimuli flashed repeatedly before it started moving, the flies followed this bar and ignored the other one, demonstrating that the flashing bar captured the flies’ attention. The study did not however, test if the flashing bar led to an increase of the perceived contrast of the subsequent stimuli, as has been seen in primates.

In this study, we therefore investigated the possibility that a cue could increase visual contrast sensitivity in insects. We conducted two different experiments with bumblebees, another model system for insect visual behaviour. In the first one, we trained bumblebees to collect a reward below a full contrast target displayed on a computer screen and tested their ability to detect the target displayed at lower contrasts when it was preceded by a cue flashed at the same location as the target, a different location or not flashed at all. Our hypothesis was that if the cue led to a spatially localized increase of their contrast sensitivity, they would be able to detect the target at lower contrast when the cue was flashed at the same location compared to the other cuing conditions.

In the second experiment, bees were trained to discriminate a full contrast target from a variable contrast distractor displayed on the opposite side of the screen. During tests, a cue could be flashed on the side of the target, on the side of the distractor or not flashed in a control condition. The prediction was that when the cue was on the side of the target, it would increase its perceived contrast and enable the bees to better discriminate it from the distractor. Conversely, when the cue was flashed on the side of the distractor, if it increased its perceived contrast, we predicted that this would hinder the bees’ ability to discriminate the target from the distractor.

## Material and Methods

### Animals and setup

We carried out the experiments on the buff-tailed bumblebee Bombus terrestris. Bumblebee colonies were purchased from commercial pollinator suppliers (Koppert BV, Netherlands and Agralan Ltd, UK) and transferred to a nest box (L=28 cm, W=16 cm, H=12 cm). The nest box had two chambers, one to house the brood and the rest of the colony and the other containing cat litter for the bees to dispose of their waste. The latter chamber was connected to a transparent tunnel leading to a foraging arena (L = 45 cm, W = 60 cm, H = 40 cm). The arena was covered with a UV transparent plexiglass board letting through the illumination from a daylight spectrum tube (Philips, Master TL5 HE 35W, 6500K) fitted to a high frequency lighting system (Philips, HF-P 1 14-35 TL5 HE III, >42KHz). The floor of the arena was covered with a random red and white checkerboard pattern to provide the bees with optic flow. The arena wall facing the tunnel exit had a computer screen (Dell S2419HGF, LCD, 1080p, 144 Hz) on which we could display visual stimuli during the experiments.

We mounted a smartphone (Huawei Nexus 6P) above the arena to record test trials at 120 fps with a 720p resolution.

The colonies had access to pollen ad libitum in a little cup placed in the nest box. During evenings and weekends, feeders filled with a 20% (w/w) sugar solution were placed in the foraging arena so the bees could feed ad libitum.

### Detection Task

This experiment tested whether flashing a cue could improve or hinder target detection by bees when the cue was on the same side or the opposite side of the computer screen relative to the target.

#### Pretraining

Individually marked bees were pretrained to collect a 50% (v/v) sugar solution reward from a little well at the centre of a transparent square plastic chip (L=2.5 cm, H=0.5 cm) placed on top of an upside-down cup (H=7 cm, D=6 cm) beneath the centre of the screen in the arena. A drop of 100 µL was placed in the well and the bee was placed over the chip with a transparent container. Once the bee started drinking, the container was removed, and the bee was allowed to return freely to her nest.

#### Training

After completing the pretraining, bees proceeded to a training phase. During this phase, we placed transparent chips on cups on each side of the computer screen. In each training bout, the screen was set to display a green background (RGB values: 0, 1, 0) and a full contrast black circular target (Diameter = 5.56 cm; RGB values: 0, 0, 0) was displayed above one of the chips. The side of the target was chosen pseudo-randomly across bouts, with a maximum of 2 consecutive bouts with the target on the same side. The chip below the target contained 100 µL of 50% Sucrose solution, while the other chip had 100 µL of distilled water. Between each trial, the chips were wiped with 70% ethanol to remove any pheromone marking and then cleaned with distilled water to remove the scent of the ethanol. We deemed a trial correct when the bee first chose the chip below the target, with a choice defined as probing the contents with her antennae or her proboscis. When a bee chose the other chip, this was deemed a wrong choice, and she was allowed to correct herself and collect the reward on the correct chip before returning to her nest. Bees were allowed to proceed to the test phase once they reached 80% success on the last 20 trials (N=18).

#### Testing

The test setup was identical to the training phase but both chips contained 100 µL of distilled water. During a test, the experimenter manually triggered the cuing as soon as the bee’s head entered the arena. This sequence involved one of three conditions. In the first, a blue square cue (side 8.33 cm) was presented for 200 ms on the side of the screen where a target would later be displayed. In the second, the cue was presented on the opposite side. In the third condition, which served as a control, no cue was presented. The position of the cue centre on the screen was 6.64 cm from the edge of the screen and from the target location (centre to centre) to prevent a masking effect (See Fig. 1). The target then appeared after a pause of 100 ms. The colour of the cue was chosen based on a previous paper demonstrating that blue stimuli disturbed shape learning in honeybees, presumably because this colour captured their attention (Morawetz et al., 2013).

**Figure 1:**
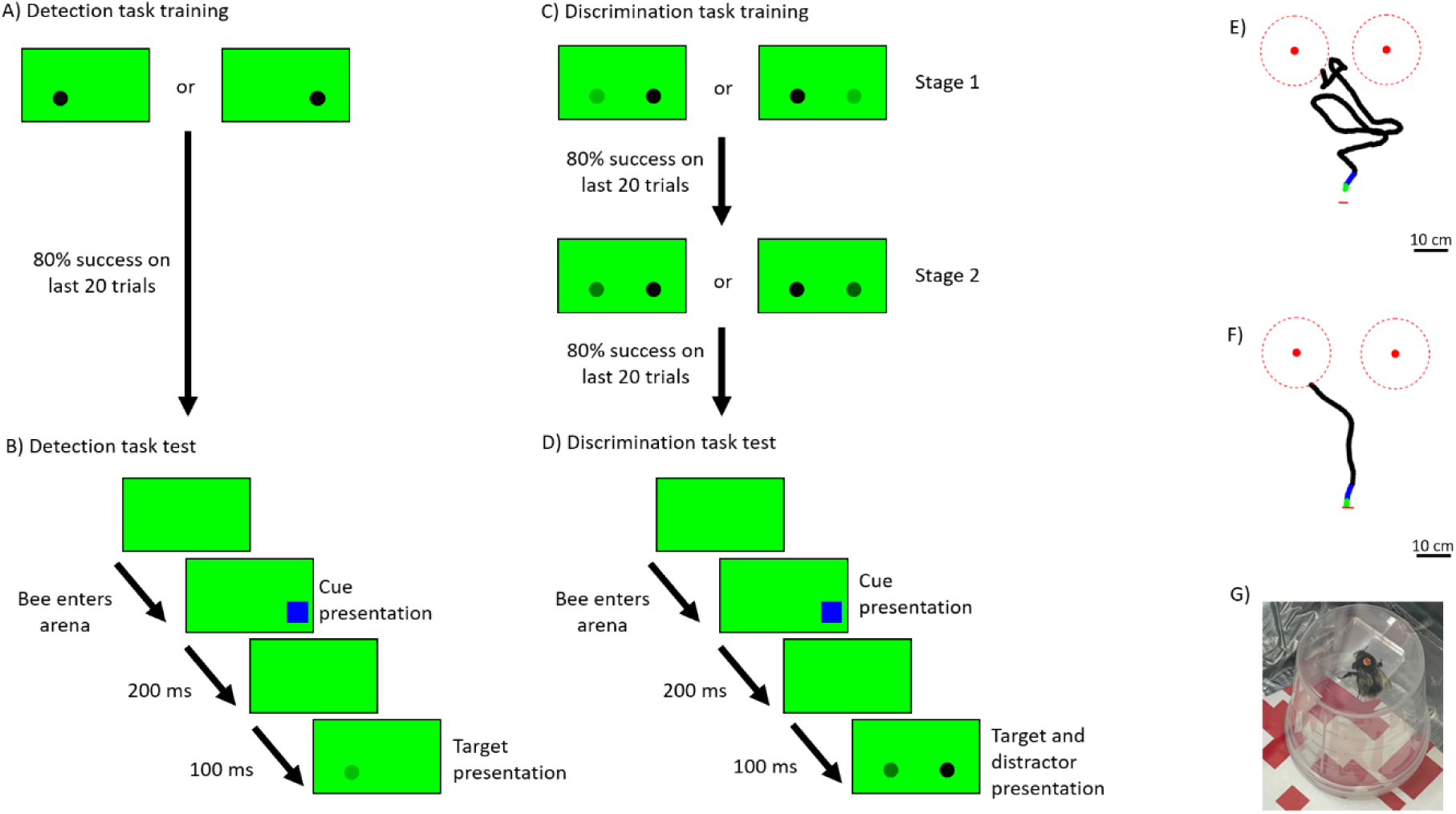
Illustration of the Detection and Discrimination experiments. A) Representations of the two possible target locations during the training phase of the detection experiment. The target was always a full contrast black circle placed either on the left or the right of the computer screen. B) Representation of a display sequence on the computer screen during a test trial of the detection task. After the bee entered the experimental arena, a blue square cue could be flashed for 200 ms on the right side or the left side of the screen or not flashed at all. After the cue disappeared, a target was presented on either side of the screen with one of 5 possible contrasts. C) Representations of the two training stages of the discrimination experiment. In stage 1, the target was always showed with a full contrast, either on the left or the right side of the screen, and the distractor was showed with a 0.448 contrast on the opposite side. Once the bee met the learning criterion, it moved to the second training stage during which the target was also showed with a full contrast, but the distractor could take one of 5 lower contrasts. D) Representation of a display sequence on the computer screen during a test trial of the discrimination task. The cuing sequence was identical to that in the detection task. During target presentation, the target was always presented at full contrast on one or the other side of the screen. On the opposite side of the screen, a distractor was displayed with one of 6 possible contrasts equal or lower to the target one. E) Top-down view of an example first approach trajectory recorded during a detection task test trial. The two red filled circles represent the two chips above which the target could appear. The dotted red circles around them represent the zones which we considered as approach zones (10 cm from the chip). The short red line shows the position of the tunnel entrance to the experimental arena. Trajectory sections up to 1 and 5 cm from the take-off point are marked in green and blue respectively. F) An example first approach trajectory recorded during a discrimination task test trial. Details as in E). G) Picture of an individually tagged bee drinking on one of the transparent chips on top of a transparent cup as used in our experiments.

Each test trial presented the bees with a target with one of 5 different contrast values (Michelson contrast: 0, 0.355, 0.615, 0.826, 1) and the side of presentation of the target was counterbalanced across trials and contrasts. Therefore, a complete test phase was composed of 30 trials (3 cuing conditions x 5 contrasts x 2 sides) which were presented in a randomised order. The test was considered finished when the bee landed on one of the chips and probed the well with its proboscis or antennae.

Between each test, we conducted two refresher bouts identical to the training bouts to keep the bee motivated. The side of the target during the first refresher was picked randomly and was counterbalanced for the second one. If a bee made a mistake during a refresher, it was repeated until she made a correct first choice.

In total, 14 out of 32 bees did not complete the training and could not proceed to the test phase either because they had died before completing the training, stopped coming out of their nest to forage or because they never reached the learning criterion and their training was abandoned. Among the 18 bees who started the test phase, 8 completed all 30 tests before dying. The sample size for each combination of cuing condition and target contrast was between 20 and 26 trials.

### Discrimination task

This experiment tested how the bee’s ability to discriminate between targets of different contrasts was affected by a cue presented on the side of the target or on the side of the distractor.

#### Pretraining

The pretraining was identical to the one of the detection task. Once the bees had learned to drink from the transparent chips, they were allowed to start the training phase.

#### Training

For the training phase, the setup was the same as the detection task. The training phase was done in two stages. In the first, we presented a full contrast black circular target (diameter = 5.56 cm, Michelson contrast = 1) on one side of the screen over one chip with 100 µL of a 50% (v/v) sugar solution. On the opposite side of the screen, we showed a similar circular distractor with a 0.448 contrast above a chip filled with 100 µL of a saturated quinine solution. The bees therefore had to learn to approach the higher contrast target regardless of which side it was displayed. After the bees had reached 80% success on the last 20 trials in the first training stage, they proceeded to the next one. In the second training stage, the target was presented at full contrast, but the distractor had a variable contrast randomly picked without replacement from 5 possible values (Michelson contrasts: 0, 0.448, 0.680, 0.826, 0.909). The list of possible contrasts was reset after the bee had experienced the 5 contrasts. Here again, a trial was marked as correct by the experimenter if the bee first chose the chip below the target with her antennae or her proboscis. Bees had to make 80% of correct choices on their last 20 trials to be selected for the test phase. Therefore, each bee experienced each distractor contrast at least 4 times during this training phase.

#### Testing

During the tests, both chips contained 100 µL of distilled water. As in the detection task, the cue consisted of a blue square (side = 8.33 cm) 6.64 cm from the edge of the screen and from the centre of the target or distractor. Three cuing conditions were implemented: the cue could be flashed on the side of the target, on the side of the distractor or not appear at all. The duration of the cue was 200 ms with a 100 ms pause before the target and distractor were displayed. The experimenter triggered the cue as soon as the bee’s head crossed the entrance of the arena. In each test trial, the target was always at full contrast while the distractor had one of 6 contrasts (Michelson contrasts: 0, 0.448, 0.680, 0.826, 0.909, 1). The side of presentation of the target and distractor were counterbalanced across trials. When the distractor had a contrast equal to the target, they were indistinguishable from one another. Therefore, to reduce the total number of trials, we presented the cue only on the side of the target in this specific distractor contrast condition (3 conditions in total, one per cuing condition). Thus, we had 33 test trials (3 cuing conditions x 6 distractor contrasts x 2 sides – 3 trials). Here again, the test was considered complete when the bee made its final choice by landing on one of the chips and probing the well with its proboscis or antennae.

Between tests, we conducted two refresher trainings, identical to the second training stage. We placed a 100 µL of a 50% (w/w) sugar reward below the target and 100 µL of quinine solution under the distractor. The target contrast was full contrast while the contrast of the distractor was one of the 5 used during the training phase. The side of presentation of each stimulus was random for the first refresher and counterbalanced for the second one. If the bee made a wrong choice, the refresher trial was repeated until she made a correct choice.

In total, 34 bees started their training and 12 finished it and moved to the test phase. Finally, 8 bees finished all tests. The sample size for each combination of cuing condition and distractor contrast varied between 10 and 20 trials.

### Video and trajectory analyses

Test trial recordings were processed with DeepLabCut™ (Nath et al., 2019) using a ResNet-50 network trained on a mix of 1377 video frames extracted from 68 trials from both experiments to analyse videos from the detection task. To analyse videos from the discrimination task, the network was trained on 1616 frames extracted from 78 flights from both experiments. This process allowed us to obtain the trajectories of bees. Since DeepLabCut made a certain number of errors, these trajectories were then cleaned from tracking anomalies using a custom-made code in R (version 4.2.3). All frames that were excluded as part of an anomaly were rebuilt by interpolation.

In order to have the most accurate take-off location for each trajectory, the videos were manually examined and the number of the frame on which the bee took off was noted. For each trajectory, if the position of the bee on the take-off frame had been interpolated, the bee coordinates were manually extracted from the video and fed back in the trajectory data.

We used the trajectories to determine which of the two chips the bee first approached in each test. This was defined as the first chip the bee approached at a distance less than 10 cm. We also measured the duration of this first approach by counting the number of video frames each bee took to complete it.

As bottom-up attention lasts for a very short time in primates (Busse et al., 2008; Hein et al., 2006; Ling & Carrasco, 2006), it was possible that the effect of the cue might be visible only at the very early stage of the flight of our bumblebees. We therefore analysed whether bees flew toward the target immediately after their take-off depending on the cuing conditions. We computed the trajectory direction of each bee relative to the target 1 cm and 5 cm away from the take-off point. To do so, we computed the angle subtended by the line joining the position of the bee when she was 1 cm (or 5 cm) from her take-off point to her take-off point and the line joining the chip below the target and the location where the bee took-off. This measures whether the direction of flight after take-off was, overall, in the direction of the target.

### Statistical analyses

Analyses were conducted using R (version 4.2.3). We analysed the results of the detection and the discrimination experiments separately but with identical methods. We ran four analyses on our data, analysing the final choices, the first approaches, the duration of the first approaches and the direction of the early trajectories.

#### Final choice

For both experiments, the final choice of the bee was recorded as 1 when the bee landed and probed the chip marked with the target and 0 when the bee probed the other chip. We then used Generalized Linear Mixed Models (GLMM, package lme4, Bates et al., 2015) to analyse this data with the choice as a dependent variable and a binomial family and a logit link function. The independent variables were the cuing condition (cue on the side of the target, cue on the opposite side, no cue), the contrast (the contrast of the target for the detection task, the contrast of the distractor for the discrimination task) and their interaction. We used the identity of the bee as a random effect.

However, for the discrimination task data, the lack of variance in the random effect (bee identity) led to a singular fit of our model. We therefore removed the random effect and used a Generalized Linear Model (GLM) to analyse this data.

#### First approach

The same analysis as above was conducted on the first approach data. If the first chip approached was the one below the target, this variable was marked as 1 otherwise it was marked as 0. A GLMM identical to the one used for the final choices was ran on this data. Here again, for the discrimination task data, the lack of variance in the random effect led to a singular fit of our model. So, we removed the random effect and ran a GLM on the data.

#### First approach duration

We also analysed the duration of this first approach with GLMMs using the glmmTMB function from the glmmTMB package (Brooks et al., 2017; McGillycuddy et al., 2025). The dependent variable was the time in seconds between the take-off and the point at which the bee was less than 10 cm away from one of the chips for the first time. In one model, we tested the effect of the contrast of the target (for the detection task) or of the distractor (for the discrimination task), the cuing condition and their interaction. While in a second model, the independent variable was the contrast, the chip chosen by the bee (correct or incorrect chip) and their interactions. These models were fitted with a Gamma family and a log link function. To improve the fit of our models used to analyse the approach duration in the detection task, the dependent variable had to be transformed by applying a log10 function to it. Finally, for both experiments, the dispersion of the data had to be modelled by providing a dispersion formula in the glmmTMB function. This dispersion was modelled based on the distractor contrast, the cuing condition and whether the chosen chip was correct or not (Dispersion formula = Contrast*Cuing condition*Chosen chip).

#### Early flight direction

We tested for the effect of the cue on the initial flight direction of the bees by analysing the bee trajectory direction relative to the target at 1 and 5 cm from their take-off point. To do so, we separated trials with the target on the right and left sides of the screen. Then, for each side group, we pooled all trials for each cuing condition and used the Rayleigh test from the “circular” R package (Agostinelli & Lund, 2024) to test whether each group’s flight direction was significantly oriented towards 0 (perfectly flying towards the target).

In addition, we used the Watson-Wheler test from the same package to test whether bee early flight direction differed between cuing conditions in each of the side groups.

### Data exclusion

We excluded flights following visual examination of the test recordings based on the following criteria: first, all trials where bees were seen crashing or landing and walking on the arena floor were excluded from all analyses (N=9 out of 335 (2.69%) for the detection task and N=1 out of 314 (0.32%) for the discrimination task). Additionally, some flights were excluded from the first approach duration and the early flight direction analyses when the bees were seen turning back to look at the tunnel entrance before approaching the screen. We believe that these sequences during which the bee faced the tunnel were learning flights and were not relevant to the target detection task (detection task: N=62 out of 326 (19.02%), discrimination task N=74 out of 313 (23.64%)). However, these flights were kept for the first approach and final choices analyses presented here. The same analyses of the bees’ choices were also run with these flights included and gave nearly identical results (not presented here).

Additionally, in the analysis of early flight direction, we only included flights in which we could see the target appear before the bee took off. These videos had either the target and/or the distractor appear on screen or we could alternatively estimate the time of target appearance based on the cue (cue appearance frame + 72 frames). For the detection task, in the uncued condition, it was often impossible to see the target if it was not at one of the two highest contrasts. Thus, we had to exclude numerous flights from the analysis for this specific condition. Finally, we only included flights in which the crossing of the 1 cm or 5 cm threshold from the take-off point happened on a frame not excluded during our cleaning and reconstruction process of the trajectories. In total, we kept 145 out of 326 flights (44.48%) for the 1 cm threshold and 146 out of 326 flights (44.79%) for the 5 cm threshold on the detection task. We also kept 92 out of 239 flights (38.49%) for the 1 cm threshold and 108 out of 239 flights (47.16%) for the 5 cm threshold for the discrimination task.

## Results

### Detection task

#### Final choice analysis

The probability of a correct final choice during tests increased with an increase in the target contrast in all three cuing conditions (Fig. 2A ; Uncued: Estimate±Standard Error=2.901±0.740, Z=3.920, p<0.001 ; Target side cued: Estimate±Standard Error=4.457±0.977, Z=4.561, p<0.001 ; Opposite side cued: Estimate±Standard Error=2.728±0.757, Z=3.605, p<0.001). However, compared to the uncued condition, the display of the cue on the target side (main effect: Estimate±Standard Error=-0.096 ±0.561, Z=-0.171, p=0.864 ; interaction with contrast: Estimate±Standard Error=1.556±1.218, Z=1.278, p=0.201) or on the opposite side of the screen (main effect: Estimate±Standard Error=0.335 ±0.550, Z=0.608, p=0.543 ; interaction with contrast: Estimate±Standard Error=-0.172±1.054, Z=-0.164, p=0.870) did not affect the chances of correct choices. There was also no significant difference between the probability of a correct final choice in the two cued conditions (main effect: Estimate±Standard Error=0.430±0.550, Z=0.782, p=0.434 ; interaction with contrast: Estimate±Standard Error=-1.729±1.232, Z=-1.403, p=0.161).

**Figure 2:**
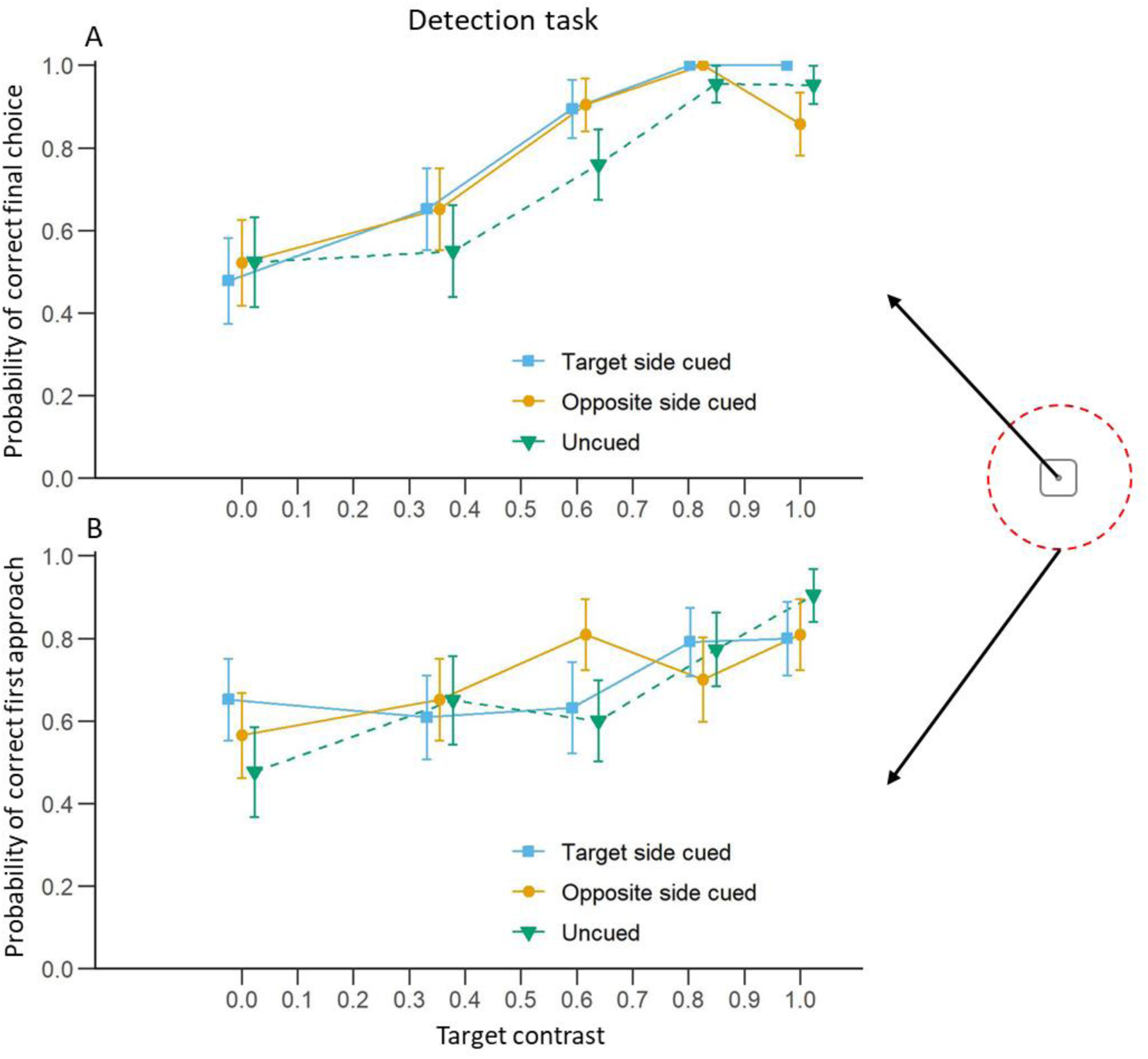
Detection task: Effect of target contrast and cuing condition on the probability of choosing the target side. A) Mean (±S.E.) proportion of trials in which bees chose target chips as a function of the Michelson contrast of the target. B) Mean (±S.E.) proportion of trials in which the bee first approached the target chip at a distance less than 10 cm. Blue curves represent trials where the cue was presented on the same side as the target. Yellow curves represent trials where the cue was presented on the opposite side to the target. Green curves represent trials without a cue. The schematic on the right shows a top view of a transparent chip. A final choice was when the bee probed the well at the centre of the chip with her antennae or her proboscis. The red dashed circle represents the 10 cm radius from the centre of the chip. A first approach was when the bee crossed the 10 cm radius around one of the two chips for the first time.

#### First approach analysis

As in the final choice analysis, an increase in target contrast increased the probability that the bees approached the correct chip in the uncued condition (Fig. 2B ; Estimate±Standard Error=1.829±0.627, Z=2.917, p=0.004). Although a similar trend was observed in the two cued conditions, the effect of the target contrast was not significant (Target side cued: Estimate±Standard Error=0.856±0.592, Z=1.146, p=0.148 ; Opposite side cued: Estimate±Standard Error=1.099±0.605, Z=1.816, p=0.069). Despite this, when compared to the uncued condition, we did not find a significant effect of the cue flashed on the side of the target (main effect: Estimate±Standard Error=0.611 ±0.530, Z=1.154, p=0.248 ; interaction with contrast: Estimate±Standard Error=-0.973±0.861, Z=-1.130, p=0.258) or on the opposite side (main effect: Estimate±Standard Error=0.532 ±0.521, Z=1.004, p=0.315 ; interaction with contrast: Estimate±Standard Error=-0.730±0.870, Z=-0.839, p=0.401) on the probability of correct first approach. In addition, the side of the cue did not have an effect on the probability of correct first approach (Target side vs Opposite side, main effect: Estimate±Standard Error=-0.080 ±0.520, Z=-0.154, p=0.878 ; interaction with contrast: Estimate±Standard Error=0.243±0.846, Z=0.287, p=0.774).

#### First approach duration analysis

We first ran a model investigating the effect of contrast and cuing condition. We found a non-significant trend that the first approach duration decreased with the increase of the target contrast when the cue was not displayed (Fig. 3A ; Estimate±Standard Error=-0.182±0.191, Z=-0.950, p=0.342). However, the same effect was significant in both conditions when the cue appeared (Target side cued: Estimate±Standard Error=-0.470±0.151, Z=-3.119, p=0.002 ; Opposite side cued: Estimate±Standard Error=-0.387±0.124, Z=-3.126, p=0.002). However, there was no significant effect of the cue when comparing both cued conditions to the uncued one (Target side cued, main effect: Estimate±Standard Error=0.136±0.170, Z=0.800, p=0.424 ; interaction with contrast: Estimate±Standard Error=-0.289±0.240, Z=-1.202, p=0.229 ; Opposite side cued, main effect: Estimate±Standard Error=0.013±0.156, Z=0.080, p=0.936 ; interaction with contrast: Estimate±Standard Error=-0.206±0.229, Z=-0.898, p=0.369). Here again, this absence of significant interaction tends to support the fact that overall, the cue did not have an effect on the first approach duration but the more the target was visible, the faster the bees made their choice. There was also no difference between the two cued conditions (main effect: Estimate±Standard Error=-0.124±0.127, Z=-0.972, p=0.331 ; interaction with contrast: Estimate±Standard Error=0.083±0.197, Z=0.423, p=0.672).

**Figure 3:**
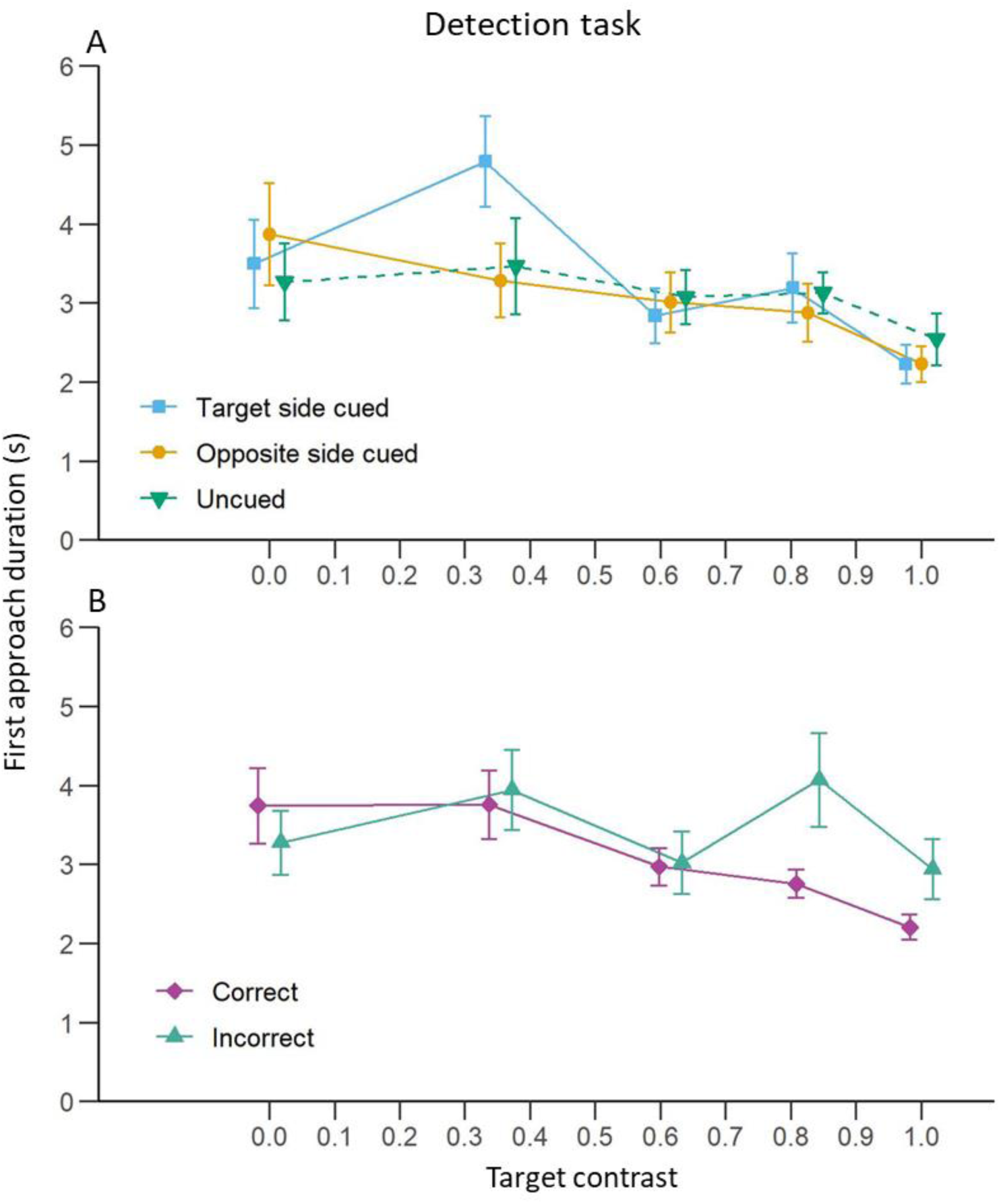
Detection task: Effect of target contrast on bee first approach duration. **A)** Mean (±S.E.) time taken by the bees to first approach either chip at less than 10 cm for each cuing condition. The blue curve represents trials where the cue was presented on the same side as the target. The yellow curve represents trials where the cue was presented on the opposite side to the target. The green curve represents trials without a cue. B) Mean (±S.E.) time taken by the bees to first approach either chip at less than 10 cm for trials where bees made a correct (purple) or incorrect (turquoise) first approach.

To investigate the effects of choice accuracy, we ran a second model with contrast and choice accuracy (correct or incorrect) as independent variables. We found a significant interaction between the two variables: bees made faster first approaches with the increase of the target contrast when they approached the correct chip (Fig. 3B ; Estimate±Standard Error=-0.477±0.100, Z=-4.781, p<0.001) but not when they approached the wrong one (Estimate±Standard Error=0.048±0.139, Z=0.344, p=0.731). This interaction indicated that although the first approach duration was similar between chips when the target was invisible, i.e. with a contrast = 0, (Estimate±Standard Error=-0.182±0.104, Z=-1.745, p=0.081), the effect of the target contrast differed depending on whether the animals were approaching the target chip or the other chip (Estimate±Standard Error=0.525±0.172, Z=3.058, p=0.002). This may indicate that the bees were more hesitant when approaching a chip when the target was not discernible.

#### Early flight direction

Bee flights in the first 1 cm from their take-off point were significantly oriented towards the target in all three cuing conditions both when the target was on the right side of the screen (Fig. 4 ; Uncued: Rayleigh test=0.901, p<0.001 ; Target side cued: Rayleigh test=0.855, p<0.001 ; Opposite side cued: Rayleigh test=0.912, p<0.001) and when it was on the left side of the screen (Uncued: Rayleigh test=0.923, p<0.001 ; Target side cued: Rayleigh test=0.953, p<0.001 ; Opposite side cued: Rayleigh test=0.960, p<0.001). The same was true for bee flights 5 cm from their take-off points when the target was on the right side of the screen (Fig. 5 ; Uncued: Rayleigh test=0.877, p<0.001 ; Target side cued: Rayleigh test=0.859, p<0.001 ; Opposite side cued: Rayleigh test=0.876, p<0.001) or the left side of the screen (Uncued: Rayleigh test=0.938, p<0.001 ; Target side cued: Rayleigh test=0.922, p<0.001 ; Opposite side cued: Rayleigh test=0.939, p<0.001).

**Figure 4:**
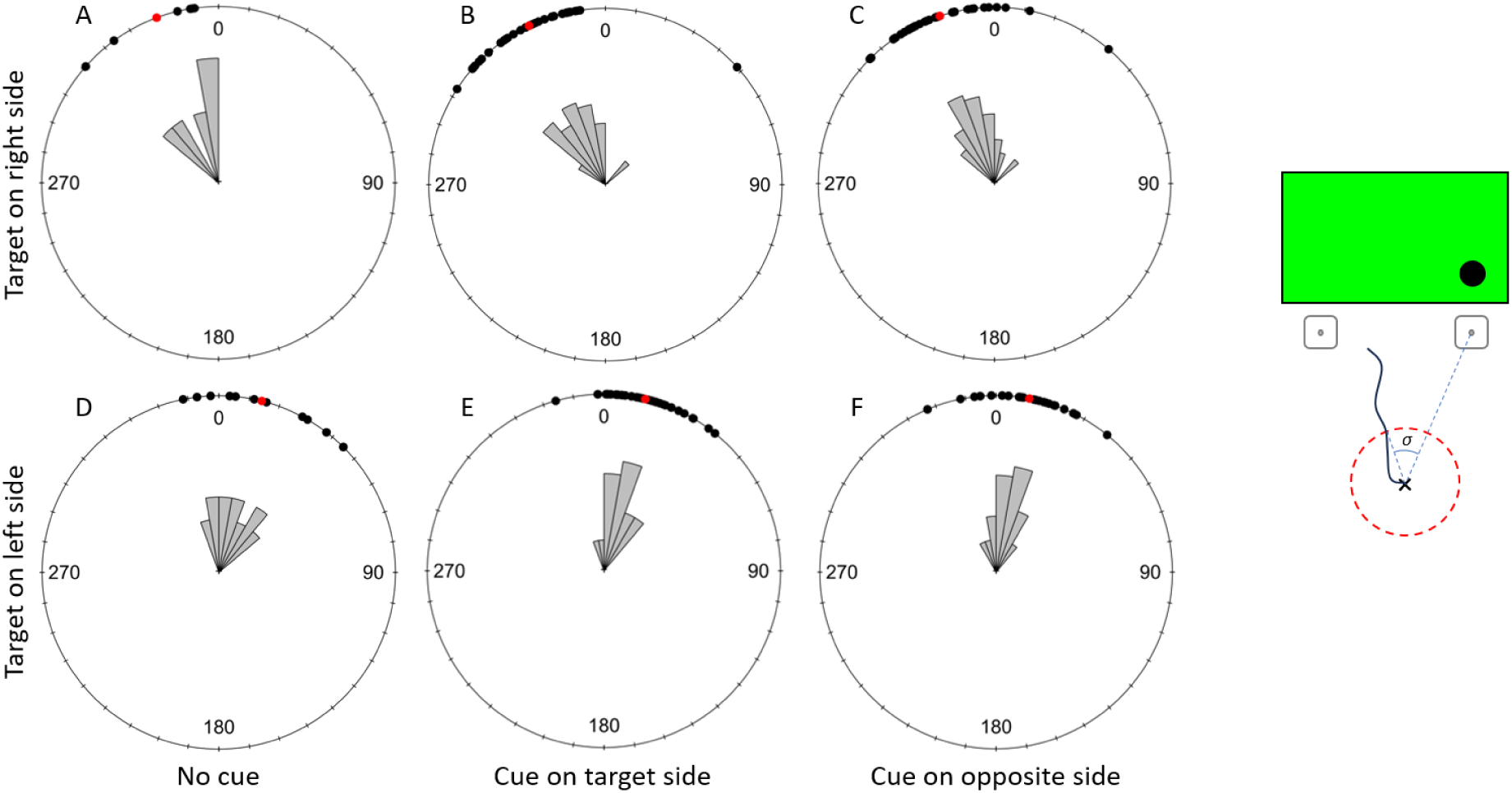
Detection task: Effect of the cuing condition on flight direction relative to the target (σ) at 1 cm from the take-off point. The top row shows trials with the target displayed on the right side of the screen and the bottom row shows trials with the target displayed on the left. A and D represent the uncued condition, B and E present data for trials with the target side cued and C and F show trials with the opposite side cued. Black dots represent individual trials, and red dots show the mean direction for each condition. 0 indicates the direction of the target. A clockwise rotation shows deviation towards the right of the target and counterclockwise indicates a deviation to the left of the target. The drawing on the right shows a schematic representation of the angle σ. It represents the two chips placed in front of the screen with the target displayed on the right side. The cross shows the bee’s take-off point, and the black line shows its trajectory. The red circle represents the early flight radius (either 1 or 5 cm in our experiment). The dashed blue lines form the angle between the place where the bee crossed the early flight radius and the direction of the correct chip. This angle therefore represents the bee’s early flight direction relative to the chip below the target.

**Figure 5:**
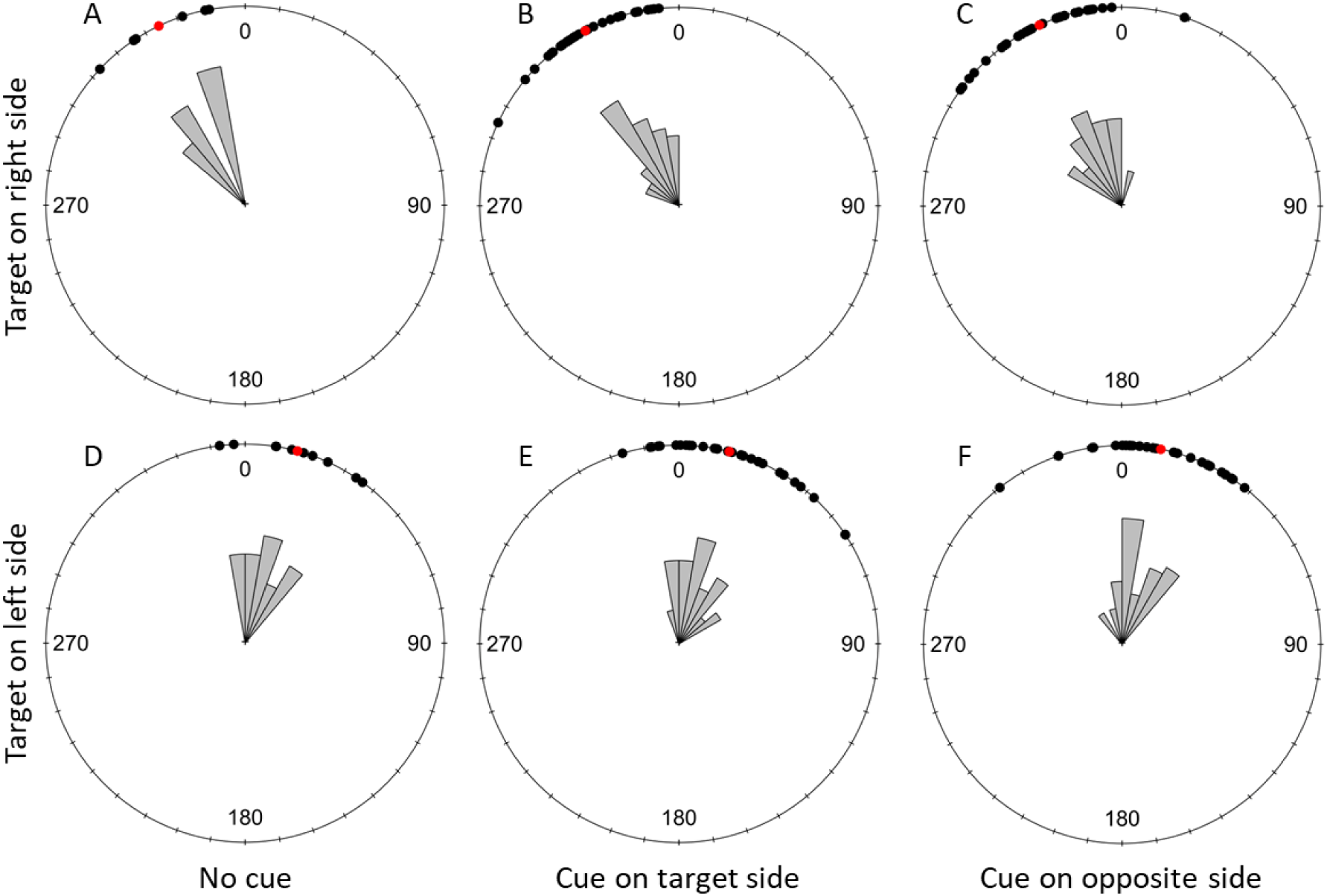
Detection task: Effect of the cuing condition on flight direction relative to the target (σ) at 5 cm from the take-off point. The top row shows trials with the target displayed on the right side of the screen and the bottom row shows trials with the target displayed on the left. A and D represent the uncued condition, B and E present data for trials with the target side cued and C and F show trials with the opposite side cued. Black dots represent individual trials, and red dots show the mean direction for each condition. 0 indicates the direction of the target. A clockwise rotation shows deviation towards the right of the target and counterclockwise indicates a deviation to the left of the target.

The direction of trajectories at 1 cm from the take-off point did not significantly differ between cuing conditions when the target was presented on the right side of the screen (W = 4.817, df = 4, p-value = 0.307) or when it was presented on the other side (W = 5.239, df = 4, p-value = 0.264). The same was true at 5 cm from the bees’ take-off point when the target was displayed on the right side of the screen (W = 2.188, df = 4, p-value = 0.701) or when it was shown on the left side of the screen (W = 0.654, df = 4, p-value = 0.957).

Because our exclusion criteria were very conservative, we ran the same analyses for the early flight directions at 1 and 5cm without excluding any flights and the results obtained were identical.

### Discrimination task

#### Final choice

In the discrimination task, increasing distractor contrast significantly decreased the probability that bees chose the target location in all cuing conditions (Fig. 6A ; Uncued: Estimate±Standard Error=-5.339±1.737, Z=-3.074, p=0.002 ; Target side cued: Estimate±Standard Error=-7.518±2.393, Z=-3.142, p=0.002 ; Opposite side cued: Estimate±Standard Error=-7.493±1.989, Z=-3.768, p<0.001). This reflects the fact that increasing distractor contrast made the target and distractor less difficult to discriminate.

**Figure 6:**
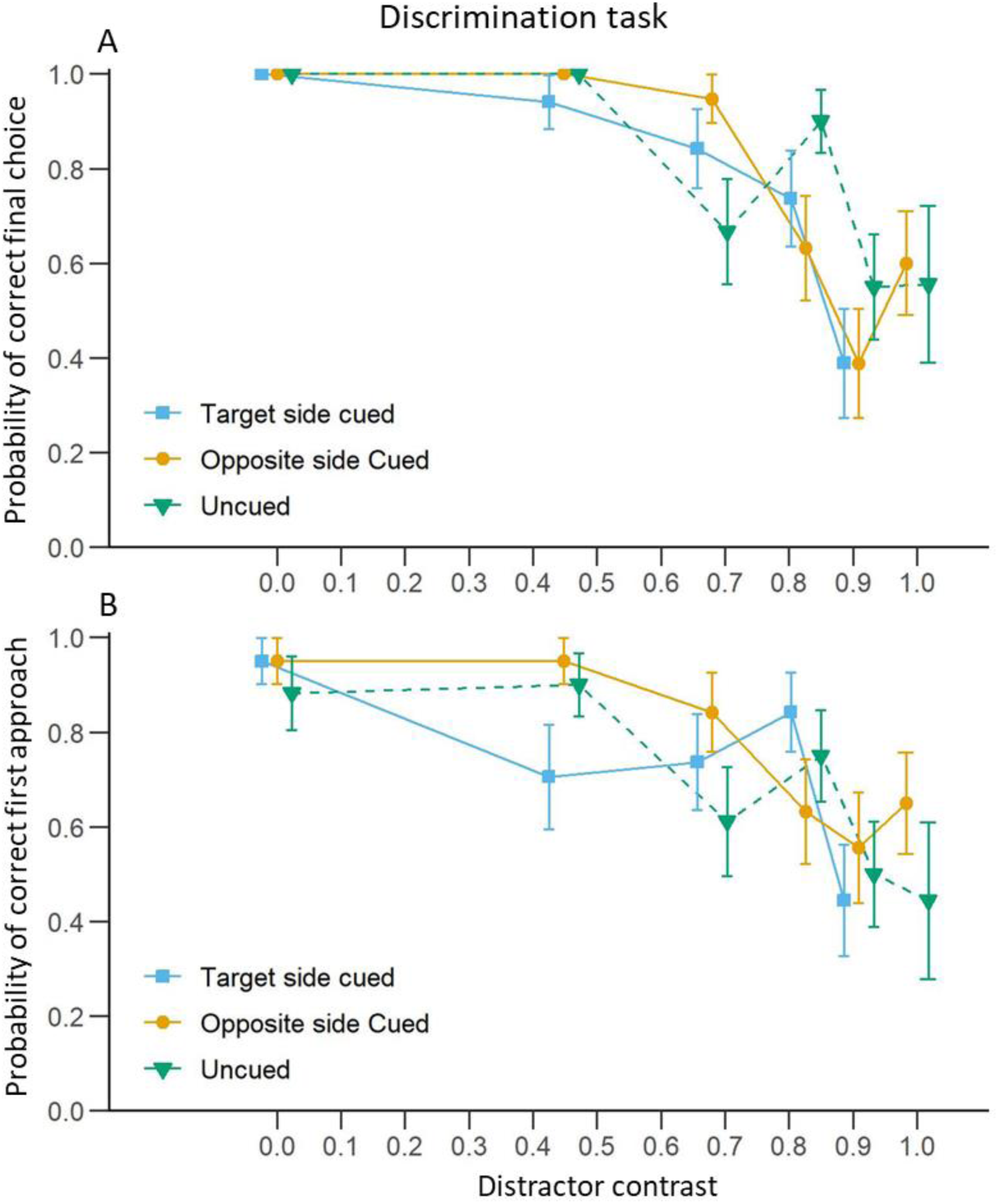
Discrimination task: Effect of distractor contrast and cuing condition on bee choices. A) Mean (±S.E.) proportion of trials in which bees chose the target chip as a function of the Michelson contrast of the distractor. B) Mean (±S.E.) proportion of trials in which bees first approached the target chip at less than 10 cm. Blue curves represent trials where the cue was presented on the same side as the target. Yellow curves represent trials where the cue was presented on the opposite side to the target. Green curves represent trials without a cue.

Cuing condition, however, did not influence the final choices of the bees when compared with the uncued condition (Target side cued, main effect: χ Estimate±Standard Error=1.372±2.440, Z=0.562, p=0.574; interaction with contrast: Estimate±Standard Error=-2.179±2.956, Z=-0.737, p=0.461 ; Opposite side cued, main effect: Estimate±Standard Error=1.772±2.288, Z=0.774, p=0.439 ; interaction with contrast: Estimate±Standard Error=-2.154±2.640, Z=-0.816, p=0.415). When comparing the two conditions where the cue was displayed, the effect of its position (on the side of the target or the side of the distractor) did not differ either (main effect: Estimate±Standard Error=0.400±2.628, Z=0.152, p=0.879 ; interaction with contrast: Estimate±Standard Error=0.025±3.111, Z=0.008, p=0.994).

#### First approach

Similar results were obtained for the probability of a first approach to the target. The bees were less likely to first approach the target side as the distractor contrast increased in the three cuing conditions (Fig. 6B ; Uncued: Estimate±Standard Error=-2.536±0.909, Z=-2.791, p=0.005 ; Target side cued: Estimate±Standard Error=-2.327±0.955, Z=-2.437, p=0.015 ; Opposite side cued: Estimate±Standard Error=-3.354±1.090, Z=-3.076, p=0.002). Here again, the cuing condition did not influence the probability of a first approach to the target when compared to the uncued condition (Target side cued, main effect: Estimate±Standard Error=-0.066±0.992, Z=-0.066, p=0.947; interaction with distractor contrast: Estimate±Standard Error=0.209±1.318, Z=0.159, p=0.874 ; Opposite side cued, main effect: Estimate±Standard Error=1.064±1.149, Z=0.926, p=0.355; interaction with distractor contrast: Estimate±Standard Error=-0.818±1.419, Z=-0.576, p=0.564). first approach probabilities in the two conditions where the cue was displayed did not differ either (main effect: Estimate±Standard Error=1.130±1.147, Z=0.985, p=0.325; interaction with distractor contrast: Estimate±Standard Error=-1.027±1.449, Z=-0.709, p=0.479).

#### First approach duration

Surprisingly, bees took less time to approach a chip as the distractor contrast increased in the uncued condition (Fig. 7A; Estimate±Standard Error=-0.206±0.096, Z=-2.152, p=0.031). This is probably explained by the fact that, contrary to what we expected, the bees were more willing to approach either chip as the distractor resembled more the target. However, this effect of distractor contrast was not observed in the two cued conditions (Target side cued: Estimate±Standard Error=-0.062±0.092, Z=-0.669, p=0.504 ; Opposite side cued: Estimate±Standard Error=-0.041±0.059, Z=-0.698, p=0.485). Despite these different of effects of the contrast in the two cued conditions and in the uncued condition, the cue did not have a significant effect on the first approach duration compared to the uncued condition (Target side cued, main effect: Estimate±Standard Error=-0.128±0.093, Z=-1.377, p=0.168 ; interaction with contrast: Estimate±Standard Error=0.144±0.133, Z=1.090, p=0.276 ; Opposite side cued, main effect: Estimate±Standard Error=-0.157±0.084, Z=-1.867, p=0.062 ; interaction with contrast: Estimate±Standard Error=0.165±0.114, Z=1.449, p=0.147). Finally, the side of the cue when it was displayed did not have an effect on the duration of the first approach (main effect: Estimate±Standard Error=-0.029±0.073, Z=-0.393, p=0.694 ; interaction with contrast: Estimate±Standard Error=0.020±0.110, Z=0.183, p=0.855).

**Figure 7:**
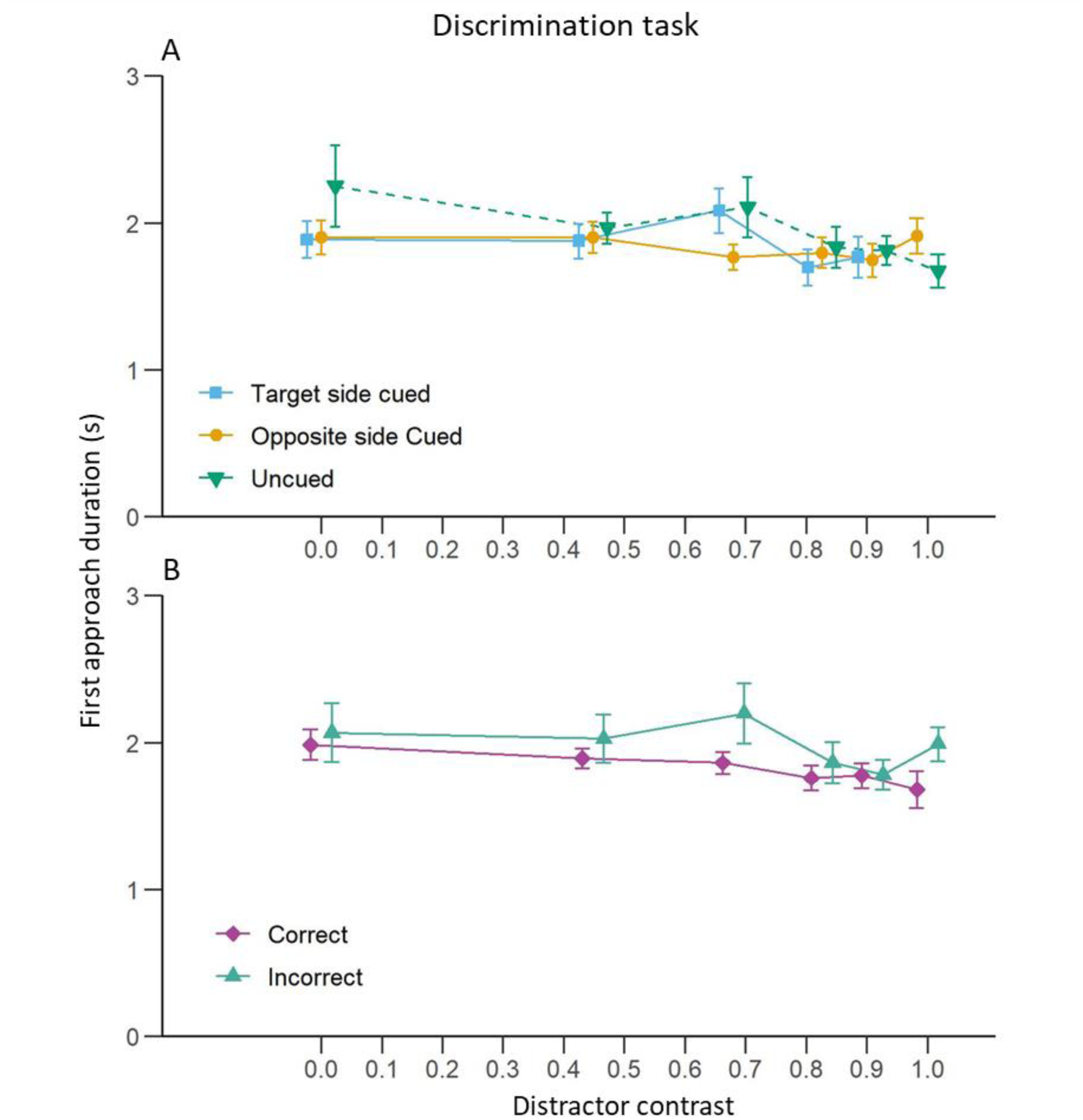
Discrimination task: Effect of distractor contrast on bee first approach duration. **A)** Mean (±S.E.) time taken by the bees to first approach either chip at less than 10 cm for each cuing condition. The blue curve represents trials where the cue was presented on the same side as the target. The yellow curve represents trials where the cue was presented on the opposite side to the target. The green curve represents trials without a cue. B) Mean (±S.E.) time taken by the bees to first approach either chip at less than 10 cm for trials where bees made a correct (purple) or incorrect (turquoise) first approach.

Contrary to our results in the detection task experiment, the distractor contrast did not significantly influence the first approach duration in trials where the bee made a correct choice (Fig. 7B ; Estimate±Standard Error=-0.087±0.048, Z=-1.825, p=0.068) but it did when the bee made a wrong choice (Estimate±Standard Error=-0.327±0.084, Z=-3.914, p<0.001) for her first approach. This was due to the fact that when bees first approached the distractor’s location and if the distractor was not visible (Michelson contrast=0), they took more time than when they approached the target (Comparing the main effect of Target side cued vs Opposite side cued conditions: Estimate±Standard Error=0.254±0.073, Z=3.484, p<0.001). However, an interaction with the contrast indicated that as the distractor became more visible, the bees approached it faster (Estimate±Standard Error=-0.240±0.098, Z=-2.441, p=0.015).

#### Early flight direction

At a distance of 1 cm away from their take-off point, bees flew significantly towards the target in every cuing condition both when the target was on the right side of the screen (Fig. 8 ; Uncued: Rayleigh test=0.918, p<0.001 ; Target side cued: Rayleigh test=0.899, p<0.001 ; Opposite side cued: Rayleigh test=0.960, p<0.001) and when it was on the left side of the screen (Uncued: Rayleigh test=0.821, p<0.001 ; Target side cued: Rayleigh test=0.743, p<0.001 ; Opposite side cued: Rayleigh test=0.798, p<0.001). The same was observed with the bees flights at 5 cm from their take-off points when the target was on the right side of the screen (Fig. 9 ; Uncued: Rayleigh test=0.936, p<0.001 ; Target side cued: Rayleigh test=0.922, p<0.001 ; Opposite side cued: Rayleigh test=0.927, p<0.001) and when the target was on the left side (Uncued: Rayleigh test=0.925, p<0.001 ; Target side cued: Rayleigh test=0.925, p<0.001 ; Opposite side cued: Rayleigh test=0.900, p<0.001).

**Figure 8:**
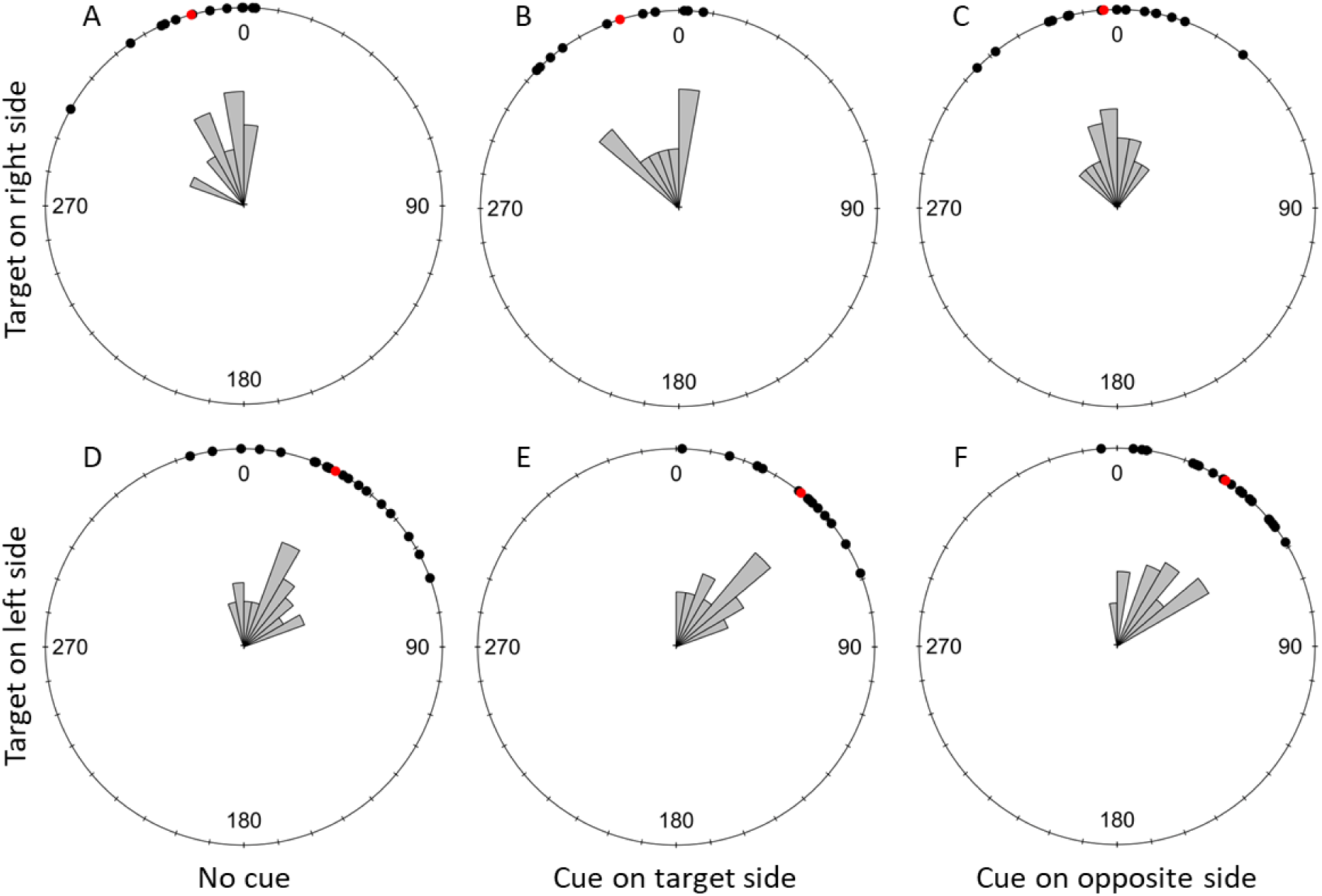
Discrimination Task: Effect of the cuing condition on flight direction relative to the target (σ) at 1 cm from the take-off point. The top row shows trials with the target displayed on the right side of the screen and the bottom row shows trials with the target displayed on the left. A and D represent the uncued condition, B and E present data for trials with the target side cued and C and F show trials with the distractor side cued. Black dots represent individual trials, and red dots show the mean direction for each condition. 0 indicates the direction of the target. A clockwise rotation shows deviation towards the right of the target and counterclockwise indicates a deviation to the left of the target.

**Figure 9:**
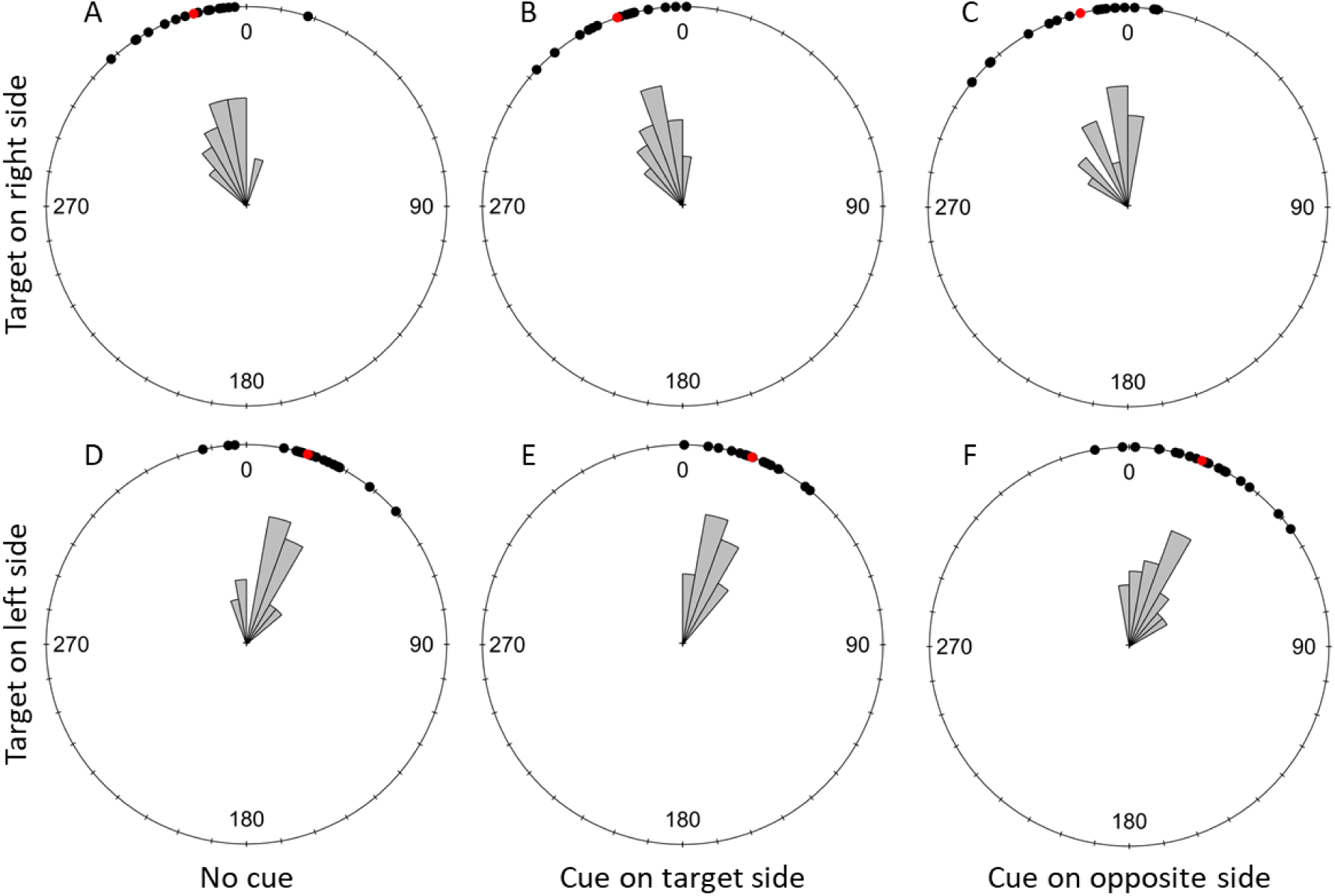
Discrimination Task: Effect of the cuing condition on flight direction relative to the target (σ) at 5 cm from the take-off point. The top row shows trials with the target displayed on the right side of the screen and the bottom row shows trials with the target displayed on the left. A and D represent the uncued condition, B and E present data for trials with the target side cued and C and F show trials with the distractor side cued. Black dots represent individual trials, and red dots show the mean direction for each condition. 0 indicates the direction of the target. A clockwise rotation shows deviation towards the right of the target and counterclockwise indicates a deviation to the left of the target.

Finally, early flight direction up to 1 cm from the take off point showed that the distribution of these did not significantly differ across cuing conditions when the target was on the right side of the screen (W = 2.288, df = 4, p-value = 0.683) or when it was on the other side of the screen (W = 3.473, df = 4, p-value = 0.482). This was also true for the flight direction at 5 cm from the take off point (target on the right: W = 5.469, df = 4, p-value = 0.243 ; target on the left: W = 2.446, df = 4, p-value = 0.654). We ran the same analyses on the flight directions at 1 and 5 cm without excluding any flights to make sure that the results presented above were not only due to our strict exclusion criteria. The analyses with the full dataset were identical.

## Discussion

The main goal of our study was to test whether a bottom-up cue could influence contrast sensitivity in bumblebees, as previous studies have shown in primates (Cameron et al., 2002; Carrasco et al., 2000, 2004; Ling & Carrasco, 2006; Z.-L. Lu & Dosher, 2000; Solomon et al., 1997). By briefly presenting a cue we attempted to capture bee attention and induce an increase of bumblebees’ contrast sensitivity in a localised area of their visual field. Our expectation was that in a detection task, if the cue was presented on the side of a subsequent target, bees would be able to detect the target at a lower contrast than if the cue was presented at another location or not flashed at all. Similarly, in a discrimination task where bees were presented a high contrast target and a variable contrast distractor, we expected the bees to have more difficulty distinguishing the two stimuli if the cue was presented on the side of a subsequent distractor.

We found that the main variables influencing the choices of the bees (both for their first approaches and final probing) was the contrast of the target in the detection task and the contrast of the distractor in the discrimination task. A previous study has shown that bumblebees in a Y-maze test can distinguish a sinusoidal grating of 0.09 cycles per degree (or 11.11°) at a Michelson contrast above 63.6% and a grating of 0.18 cycles per degree (or 5.56°) at contrasts above 81% (Chakravarthi et al., 2016). Because our target and distractor subtended a visual angle of around 6.8° from the tunnel entrance, we expected that at the lowest contrasts, the stimuli would not be detectable by the bees. Therefore, in the detection task, the bees were more likely to make a random choice at these contrasts. Conversely, during the discrimination task, as the contrast of the distractor increased it became less distinguishable from the full contrast target and thus the discrimination between the two became more difficult for the bees.

Target contrast also influenced first approach duration during the detection task. When the target was barely visible or altogether absent, bees took longer to first approach a chip compared to when the target was clearly visible. This result indicates that as a result of the successful training, bees were really looking for the target and were reluctant to approach a chip when they could not see one, even without a punishment for wrong choice. The contrast of the distractor in the discrimination task, however, had the opposite effect on first approach duration. This was surprising because, as the contrast of the distractor increased, it was more similar to the full contrast target. Discriminating between the two therefore became more difficult (as confirmed by the final choices and first approach probabilities of the bees). Thus, we could have expected bees to show the same increased approach duration as we saw in the detection task when the target was less visible, making the choice more difficult. Instead, the bees appear to simply choose one chip (right or wrong) and fly directly towards it, even though a wrong choice was punished by the taste of quinine during the discrimination task training. This suggests that our bees did not really compare the target and distractor contrasts but only estimated whether the one they first detected was close enough to the full contrast and, if so, approached and landed on the chip bellow it.

We expected early flight orientations to be random when the target was invisible in the detection task and to grow more oriented towards the target as its contrast increased. For the discrimination task, we expected bees to be oriented early on towards the target when the distractor was invisible and to get progressively more randomly oriented as the distractor contrast increased and was less distinguishable from the target. Contrary to these expectations, target contrast did not influence the early flight orientation of bees in the detection task, and the same was true for distractor contrast in the discrimination task. One theoretical explanation for this could be that the bees could not see the target from the entrance of the tunnel. However, our target subtended a visual angle of approximately 6.8° at the entrance and bumblebees can resolve achromatic sinusoidal gratings of 0.21 cycles per degree or 4.76° (Chakravarthi et al., 2016). Other research also shows that bumblebees can detect yellow targets sustaining angles between 3.4° and 7° depending on the size of the bee (Spaethe & Chittka, 2003), with only the smallest individuals needing target larger than 6°. Moreover, later studies have demonstrated in Y-maze experiments that bumblebees could detect similar yellow targets subtending an angle as small as 2.3° (Dyer et al., 2008) or 1.8° (Wertlen et al., 2008). Thus, it seems unlikely that the bees were unable to resolve the target at the tunnel entrance. It is perhaps more likely that they made their decision at a later point when they were closer to the screen.

More importantly, we didn’t see any effect of the cue on the bees’ ability to detect the target or discriminate it from the distractor. One possibility is that the bees did not perceive the cue either due to its size or to its duration. However, this seems unlikely. The measured irradiance of the cue against the background provides a strong chromatic and achromatic contrast (achromatic Michelson contrast=0.83). The cue also subtends an angle of around 9.7° at the tunnel entrance which bees would be able to resolve (Chakravarthi et al., 2016; Dyer et al., 2008; Spaethe & Chittka, 2003; Wertlen et al., 2008). It is also unlikely that the duration of the cue was too short for the bees to perceive it. The integration time of Bombus terrestris’ blue photoreceptors is 9.7 ms (Skorupski & Chittka, 2010), well below the duration of our cue. In addition, this species of bumblebee was behaviourally shown to detect blue bars flashed for as short as 25 ms (Nityananda et al., 2014). Thus, the bees should have been able to perceive our cue with a presentation duration of 200 ms.

Flashing cues have also been shown capture insect attention (Sareen et al., 2011). Fruit flies were more likely to follow a vertical bar on a circular screen if it flashed multiple times before to moving. Our cuing paradigm does differ from the one used in this experiment. There, the target was identical to the cue and flashed repeatedly at 10 Hz, while in our case, the cue was presented only once for a duration of 200 ms rather than flashing on and off. Our cue was also distinct from and did not spatially overlap with the target location to avoid a possible masking effect. Similarly to Sareen and colleagues, other experiments have also used cues that are identical to the targets and showed attentional capture. Lancer *et al*. (2019) recorded from the CSTMD1 neuron of dragonflies in response to two targets simultaneously moving upward. They showed that the neuron had equal chances to selectively attend to either one of the targets. However, if one target appeared earlier than the second one both temporally and spatially on the screen, it was more likely to be attended by the neuron. Here again, the cue and the target were identical. Finally, the bees in our experiment were freely flying compared to the tethered flies and dragonflies in the previous research. We were trying to better recreate classic spatial cuing experiments with our paradigm but this appears to not have been effective in capturing attention. If attentional processes in insects are comparable to those of vertebrates, the nature of the cue, its position relative to the target or the time between the cue and target onset would be important to successfully capture the animal’s attention (Franconeri et al., 2005; Franconeri & Simons, 2003; Fuller et al., 2009; S. Lu, 2006; Posner & Cohen, 1984; Pratt & McAuliffe, 2001; Steinman et al., 1997; Tsal, 1983). Our results suggest that our cue did not possess the required characteristics or duration to capture bumblebees’ attention.

Since worker bumblebees search and forage on rewarding flowers in their environment (Heinrich, 1976), it might be more critical for them to evolve a cognitive process resembling vertebrate top-down attention. Such a process would allow them to tune their sensory system to enhance the detection of stimuli associated with the most rewarding flowers in their environment such as their shape or colour (Liu, 2019). On the other hand, bumblebees are predated by birds at the entrance of their nest (Goulson et al., 2018) or by wasps (Dukas, 2005) and crab spiders while foraging on flowers (Dukas & Morse, 2003; Morse, 1986; Rodríguez-Gironés & Jiménez, 2019). Crab spiders are sit-and-wait predators that are camouflaged on flowers and strike at bumblebees after they land to forage. To avoid being predated by these animals, it would be beneficial to have the ability to prioritise sudden changes in the environment (such as a predator attack). Having a faster reaction time as a result would be essential for the workers’ survival. As the predation on workers decreases bumblebee colony fitness (Goulson et al., 2018), with an especially strong effect of crab spiders’ predation (Cresswell, 2017), we should expect some form of bottom-up attention in bumblebees to help them evade attacks. Given our results, investigating this would likely need different approaches to the one we took.

Recently, studies have shown that fruit flies have dedicated neural pathways responding to visual looming cues and their characteristics such as size and direction to generate fast and directed non-stereotypical escape behaviours, overriding other behaviours (Ache et al., 2019; Card & Dickinson, 2008; de Vries & Clandinin, 2012). Such a rapid and spontaneous analysis of a stimulus strongly resembles bottom-up attention. This suggests that looming cues may be better suited to investigate similar attention-like processes in insects.

## Author Contributions

VN obtained the supporting fundings. VN and TR designed the experiment. MC, CS and TR conducted the experiment. TR analysed the videos. TR ran the statistical analyses and wrote the paper. VN Edited the manuscript.

## Acknowledgements

VN and TR are supported by a BBSRC David Phillips fellowship BB/S009760/1 to VN.

## Declaration of Interests

The authors declare no competing interests.

## Notes

### Competing Interest Statement

The authors have declared no competing interest.

