## Supplementary Information and Figures for "Testing bottom-up cuing effects on target detection and discrimination in Bumblebees"

<u>Description of the post-processing of the DeepLabCut™ data:</u>

All the post processing of the trajectory data was made in R (version 4.2.3).

The DeepLabCut™ neural network was trained to detect the head, thorax, and the tip of the abdomen of bees to compute their orientation. In addition, it was also trained to detect the centre of the two transparent chips, the bottom corners of the squared shaped tunnel entrance and two screws on each side of the arena that were used as reference points. For each test trial video, we used these two points to standardise the coordinate plane: the left screw when facing the screen was set at coordinate X=0 cm, Y=0 cm and the second screw was set at coordinates X=0 cm and Y=59.50 cm. Thus, from the coordinates of these two points given in pixels by DeepLabCut™, we could compute the ratio of pixels per centimetre for each video as well as an angle of deviation from the horizontal axis. With these two pieces of information, we converted each pixel coordinate given by DeepLabCut™ for every point to centimetres and rotated the plane of coordinates to make the axis between the two reference points the horizontal axis.

<u>*Confidence level and boundaries filter:*</u> The tracking of bee heads and abdomens were not very accurate and we could not use these points to compute the orientations. The tracking of the thorax was more accurate and we used this to obtain the bees' positions and rebuild their trajectories. Because there were still many instances of tracking errors, this data was submitted to various steps of filtering. First, all frames where the thorax was out of the edges of the arena were discarded (X limits: -20 cm, 80 cm ; Y limits: -30 cm, 30 cm). We also excluded all frames where DeepLabCut™ gave a confidence level lower than 0.98 for the detection task and 0.95 for the discrimination task.

<u>*Surrounding frames distances filter:*</u> We next implemented a second level of filtering designed to detect isolated or small groups of frames where the tracking of the position of the bee was wrong and deviated significantly from the trajectory (Fig. S1A). We would expect the bee positions on these frames to be farther away from the trajectory than any other given frame, leading to an increase in the median distance to all the other frames. To capture this difference, for each frame (hereafter "the focal frame"), we computed the distance between the bee's position on this focal frame and the position in each of the 60 frames before and each of the 60 frames after. Where 60 frames before or after were not available, such as at the beginning or the end of the video, distances were calculated for the frames available. This gave us a distribution of 120 distances from the focal frame to the other frames on the trajectory.

To obtain the control distribution of distances within a trajectory, we calculated distances between the bee positions in the 60 frames before and the 60 frames after two by two, starting from the most extreme ones (the first frame of the 60 before vs the last of the 60 after the focal one), finishing with the two frames immediately surrounding the focal frame.

After obtaining these two distributions of distances, we computed their respective medians and the ratio of the medians:

Median distances from all other frames to the focal frame/Median distance of all other frames to each other.

If this ratio was above 1.1, the focal frame was deemed a deviation from the real trajectory and was discarded. We relied on computing the medians since we were dealing with groups of tracking errors rather than a single deviation.

*Freeze detection filter:* A third level of filtering was then applied to the trajectory data. This filtering step was designed to address short freezes when recording the videos. During these freezes, subsequent frames were almost identical. Therefore, DeepLabCut™ gave very similar coordinates in consecutive frames. To detect these freezes, we computed the distance between the bee position between each consecutive frame. If this distance was lower than 0.0001 cm, this frame was then discarded. We then checked the distance between the first frame and each of the frames following the discarded frame. We discarded these until we found frames with distances to the first frame greater than 0.015 cm. This distance was chosen because it would compare to the typical distance travelled between frames by a slow walking bumblebee.

*Frame to frame distance filter:* The fourth process of filtering aimed to catch tracking errors that were missed by the earlier levels of filtering. This could happen when the tracking errors alternated with correct tracking points, such that the tracked location was almost identical when an error was made or if there was a long succession of errors (>60 frames). In these situations, the second level of filtering would not be able to discriminate tracking errors from the real trajectory because the median distances as computed in that filtering would be very similar.

We therefore implemented an additional filtering (Fig. S1B). For each trial, we selected all the frames that were not discarded by the previous filtering steps and marked the first one as “good”. We then computed the distance between the bee position on the next non-discarded frame to that of the “good” one. We then divided this distance by the number of actual frames (including discarded frames) separating these two frames. If this ratio was more than 2 cm/frame, the focal frame was marked as “bad”. This was to identify significant deviations from the trajectory.

For the next frame, the distances/frames were computed from both the last known “good” frame and the last known “bad” frame. If the distance/frames ratio was large (more than 2 cm/frame) from the last “good” frame and small (less than 2 cm/frame) from the last “bad” frame, the treated frame was marked as “bad”. This was to identify frames following a deviation where DeepLabCut continued to wrongly identify the bee in positions close to the deviation.

If the ratio from the last “good” frame was small, and the ratio from the last “bad” frame was large, the treated frame was marked as “good”. This would represent a point where DeepLabCut now correctly identified the bee on the trajectory.

If the ratio from the last “good” frame and from the last “bad” frame were both large, the treated frame was marked as “good”. This would also represent a point where DeepLabCut now correctly identified the bee on the trajectory after long sequence of tracking errors.

Finally, if the ratio from both the last “bad” frame and the last “good” frame was small, this could represent an even longer sequence of misidentified positions such that the number of frames to the good frame increased enough to reduce the ratio to that frame significantly. In this case, we checked the raw distance from the last “bad” frame. If it was less than 2 cm, we marked the frame as “bad”. Otherwise, we considered that frame as “good”. All “bad” frames were then discarded from the trajectory dataset.

*Rebuilding of trajectory:* To obtain the bees’ take-off position for our orientation analyses, we first checked the videos and noted the frame number on which the bee took off. We then checked whether this frame was discarded from the trajectory obtained with DeepLabCut™. If the frame was discarded, we visually obtained the bee position directly from the video recording and corrected the coordinates for this frame in the trajectory.

Finally, the bee position on all discarded frames was rebuilt by linear interpolation.

At the end of this process, we plotted and visually checked all trajectories and we were eventually satisfied that our filtering process had removed all errors in the tracking of the bee positions (Fig. S2).

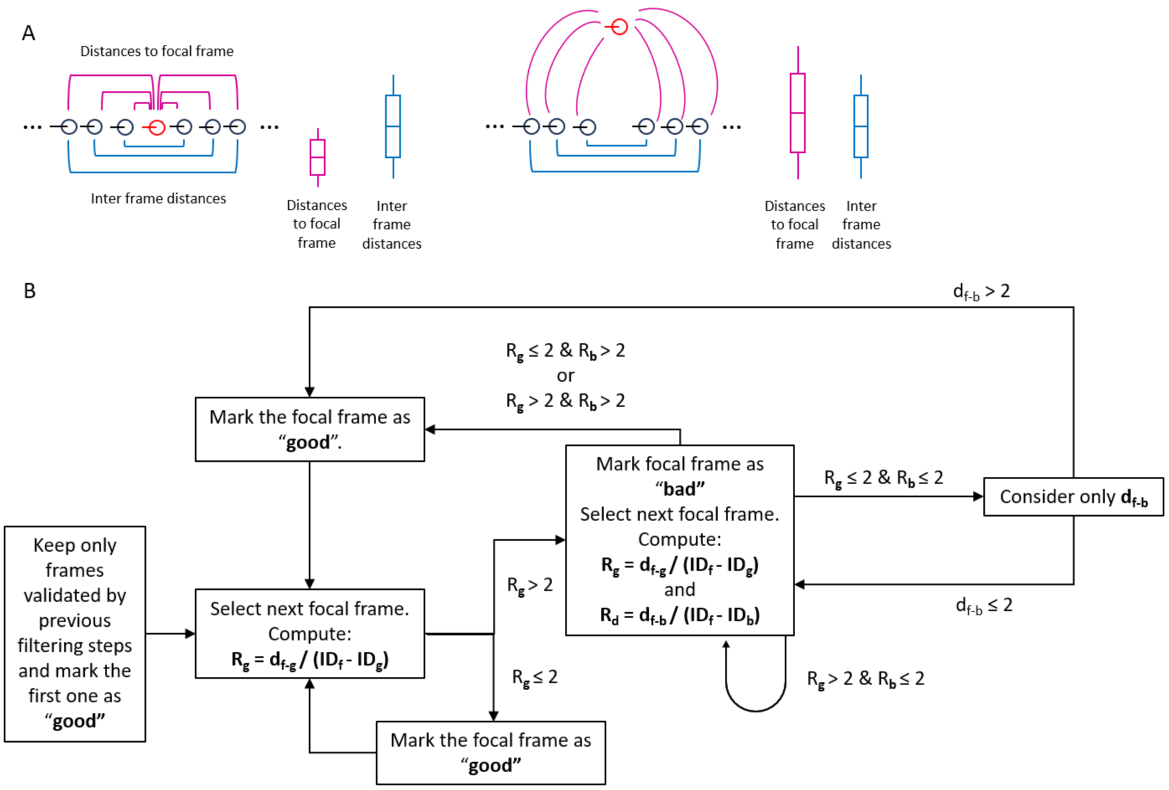


$d_{f\text{-}g}$: Euclidian distance between the bee positions on the focal frame and on the last frame marked as "good".
$d_{f\text{-}b}$: Euclidian distance between the bee positions on the focal frame and on the last frame marked as "bad".
$ID_f$: Frame number in the video for the focal frame.
$ID_g$: Frame number in the video for the last frame marked as "good".
$ID_b$: Frame number in the video for the last frame marked as "bad".

**Figure S1: Illustration of the filtering process during the post-processing of the DeepLabCut™ data.** A) Schematics showing the logic behind the *surrounding frames distances filtering* process. The circles represent the positions of a bee on the successive video frames. In red, is the focal frame (the frame currently being treated to determine if the bee position is correctly tracked by DeepLabCut™ or it if is an anomaly in the bee trajectory). During this filtering process, we computed the distance between the bee position on the focal frame and on the frames before and after (represented in pink). We then computed the distances between the bee positions on the frames surrounding the focal one, two by two (as represented in blue). If the focal frame is rightfully part of the bee trajectory (as shown in the left schematic), the distances between the bee position on the focal frame and the other frames should usually be shorter than the distances between the bee positions on the surrounding frames between themselves (as illustrated by the two boxplots on the left schematic). Alternatively, if the bee position on the focal frame is wrong, it should be offset from the real trajectory (schematic on the right) and thus the relation between the two distributions of distances should be inverted (as illustrated by the boxplots on the right). Therefore, comparing the medians of the two distributions is a good indication of the validity of the bee position on the focal frame. B) Flow chart showing the *frame to frame distance filtering* process. For each video trial, we removed all frames discarded by the previous filtering steps. Then, the current process was recursively applied to each of the remaining frames to determine whether the bee's position on them was accurately tracked by DeepLabCut™ (marked as ”good”) or not (marked as ”bad”). To do so, we examined whether the detected bee on the focal frame travelled a reasonable distance (with a threshold of 2 cm per frame) from its position on the last known ”good” and last known ”bad” tracking frames. All framed marked as ”bad” were eventually discarded.

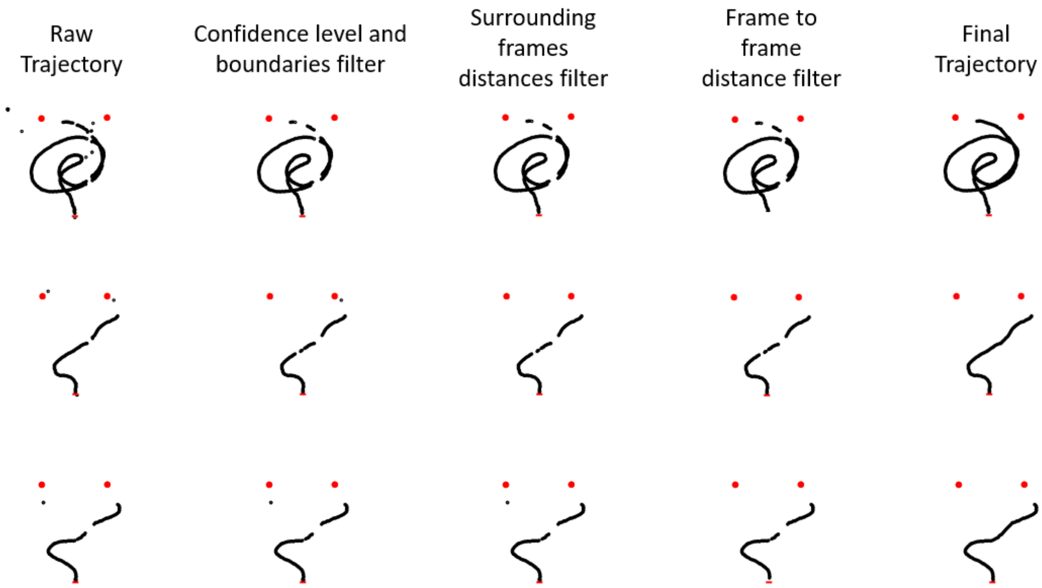


**Figure S2: Three examples of top-down view of first approach trajectories at different stages of the post-processing.** From left to right are the raw data from DeepLabCut™ standardised but not filtered, the trajectories after the *Confidence level and boundaries filter*, the *surrounding frames distances filter* and the *frame to frame distance filter*, and finally the final trajectories with the gaps interpolated. The red circles represent the position of the two artificial flowers, the red bar shows the location of the tunnel entrance, and the black circles represent the position where the bee was detected on each frame. Tracking anomalies can be seen as isolated groups of points separated from the rest of the trajectories. For numerous trajectories, the filtering based on the confidence level given by DeepLabCut™ was sufficient to remove all anomalies (as seen in the example on the top row). However, some anomalies were marked with a high confidence level by DeepLabCut™. These could be detected and removed by the *surrounding frames distances filter* step (middle row) or the *frame to frame distance filter* one (bottom row).
